# Examining the neural antecedents of tics in Tourette syndrome using electroencephalography

**DOI:** 10.1101/2020.05.01.071837

**Authors:** Barbara Morera Maiquez, Georgina M. Jackson, Stephen R. Jackson

## Abstract

Tourette syndrome (TS) is a neurological disorder of childhood onset that is characterised by the occurrence of motor and vocal tics. TS is associated with cortical-striatal-thalamic-cortical circuit [CSTC] dysfunction and hyper-excitability of cortical limbic and motor regions that are thought to lead to the occurrence of tics. Importantly, individuals with TS often report that their tics are preceded by ‘premonitory sensory/urge phenomena’ (PU) that are described as uncomfortable bodily sensations that precede the execution of a tic and are experienced as a strong urge for motor discharge. While tics are most often referred to as involuntary movements, it has been argued that tics should be viewed as voluntary movements that are executed in response to the presence of PU and bring temporary relief from the uncomfortable bodily sensations that are associated with PU. This issue remains unresolved but has very important implications for the design of clinical interventions for TS. To investigate this issue further, we conducted a study using electroencephalography (EEG). Specifically, we recorded movement-related EEG (mu and beta band oscillations) during (a) the immediate period leading up to the execution of voluntary movements by a group of individuals with TS and a group of matched healthy control participants, and (b) the immediate period leading up to the execution of a tic in a group of individuals with TS. We demonstrate that movement-related mu and beta band oscillations are *not* observed prior to tics in individuals with TS. We interpret this effect as reflecting the greater involvement of a network of brain areas, including the insular and cingulate cortices, basal ganglia nuclei, and the cerebellum, in the generation of tics in TS. We also show that beta-band desynchronization does occur when individuals with TS initiate voluntary movements, but, in contrast to healthy controls, desynchronization of mu-band oscillations is not observed during the execution of voluntary movements for individuals with TS. We interpret this finding as reflecting a dysfunction of physiological inhibition in TS, thereby contributing to an impaired ability to suppress neuronal populations that may compete with movement preparation processes.

## Introduction

Tourette syndrome (TS) is a neurological disorder of childhood onset that is characterised by the presence of chronic vocal and motor tics (Cohen, Leckman, & Bloch, 2013). Tics are repetitive, stereotyped behaviours that occur with a limited duration (Cohen et al., 2013). Motor tics can be simple or complex in appearance, ranging from repetitive movements to coordinated action sequences. Verbal tics can consist of repetitive sounds, words or utterances, the production of inappropriate or obscene utterances, or the repetition of another’s words. Tics occur in bouts, typically many times in a single day, and are the most common form of movement disorder in children.

The volitional nature of tics has been a topic of debate (Cavanna, Black, Hallett, Voon, 2017). While many refer to tics as involuntary movements (e.g., Ganos, Asmuss, Bongert, Brandt, Münchau, Haggard, 2015), others have argued that tics are instead voluntary, and occur in response to so-called ‘premonitory urges’ (Leckman, Bloch, Scahill, King, 2006). Thus, the majority of individuals with TS report that their tics are often preceded by premonitory sensory/urge phenomena (PU) that are described as uncomfortable cognitive or bodily sensations that occur prior to the execution of a tic and are experienced as a strong urge for motor discharge (Cohen et al., 2013). Individuals who experience PU often report that: these experiences are more bothersome than their tics; that expressing their tics give them relief from, and temporarily *abolishes*, their PU; and that they would not exhibit tics if they did not experience PU. For this reason, it has been proposed that PU should be considered as the driving force behind the occurrence of tics, and that tics are a learnt response to the experience of PU (Leckman et al., 2006; Cavanna et al., 2017).

One important argument for distinguishing tics from volitional movements is the finding that motor tics do not exhibit the peri-movement electrical (EEG) potentials that are observed immediately before volitional movements (Obeso, Rothwell, Marsden, 1981). Specifically, voluntary movements are typically associated with distinct pre-movement ‘readiness’ (RP), a motor (MP), and post-movement potentials. The pre-movement (RP) potential is represented by a negative wave beginning about 1.5 or 2 seconds before the movement which leads to a rapid rise in negativity at around 500ms before movement onset. The MP is represented by a positive peak and occurs with movement onset and is localized to the contralateral primary sensorimotor cortex (SMC). Previous studies have shown that while voluntary movements produced by individuals with TS are preceded by the normal peri-movement electrical potentials as observed in typically developing individuals (Karp, Porter, Toro, & Hallett, 1996; Obeso & Marsden, 1981; Van Der Salm, Tijssen, Koelman, & Van Rootselaar, 2012), these potentials are reported to be absent prior to, and at the time of, motor tics (Obeso et al., 1981). Although a subsequent study reported that tic generation was associated with an RP in 2 out of 5 patients (Karp et al., 1996).

Volitional movements also lead to robust modulation of movement-related brain oscillations in the mu (8-12Hz) and beta (13-30Hz) frequency bands. Specifically, there is a substantial decrease in power (event-related desynchronization - ERD) in both frequency bands that typically commences shortly before movement onset, and is sustained for the period of movement execution. After the movement is terminated, there is a substantial increase in power (event-related synchronisation - ERS) for both rhythms. The ERD is thought to reflect cortical activation, while ERS is thought to reflect reduced cortical excitability or inhibition. This is supported by TMS studies, which have shown that there is an increase in motor excitability just prior to a movement (80-100ms before movement) (Chen et al., 1998), during a movement (Chen et al., 1998); and that motor cortex excitability then decreases ~500-1000ms after the movement has completed (Chen, Corwell, & Hallett, 1999). For beta-band oscillations this is often referred to as the post-movement beta rebound (PMBR). It has been suggested that these movement-related brain oscillations, particularly the beta-band oscillations, reflect an internal estimate of the likelihood of the need for a novel voluntary action, and in effect signal motor readiness (Jenkinson and Brown, 2011). If this is the case then one might make the following predictions. If tics are indeed *voluntary* movements executed intentionally to relieve PU, then we would expect to see clear evidence for movement-related ERD when individuals with TS execute volitional movements, and also when they tic. By contrast, if tics are essentially *involuntary* movements, then we would expect to see clear evidence for movement-related ERD when individuals with TS execute volitional movements, but not necessarily when they execute motor tics. The current study investigated this issue using EEG-recording techniques.

## Methods: Study 1. Neural antecedents of volitional movement

### Participants

Seventeen individuals (7 females; aged between 12 and 51 years; mean age of 23 years) with a confirmed clinical diagnosis of TS were recruited to this study. Three participants were left-handed. Participants with co-occurring neuropsychiatric conditions were not excluded. Data from two TS participants was excluded due to excessive noise caused by the frequent and intense tics during EEG recording. The remaining 15 participants included 7 females aged between 12 and 35 years (mean age = 20.3 years). Three participants were left-handed. Details from TS participants can be found in Table 1. Fifteen individuals (9 females; aged between 19 and 28 years; mean age of 22.5 years) were recruited as controls. One participant was left-handed. All participants gave informed consent and the study was approved by a University of Nottingham ethical review committee.

**Table 1.** Characteristics of TS participants in Study 1

| TS ID | Age | Gender | Handedness | Medication | Comorbidity | WASI | PUTS (max32) | Impairment | Motor | Vocal | Global |
| --- | --- | --- | --- | --- | --- | --- | --- | --- | --- | --- | --- |
| TS184 | 19.37 | M | RH | Sulpiride 400mg | None | 82 | 30 | 50 | 21 | 21 | 92 |
| TS212 | 15.44 | F | LH | None | None | 110 | 29 | 49 | 19 | 13 | 81 |
| TS182 | 18.25 | F | LH | Clonidine 4x25micrograms/day | None | 85 | 18 | 20 | 21 | 9 | 50 |
| TS213 | 21.39 | F | RH | Citalopram 20mg/day | None | 109 | 22 | 45 | 22 | 16 | 83 |
| TS030 | 22.34 | M | RH | None | None | 103 | 7 | 17 | 15 | 8 | 40 |
| TS204 | 33.19 | M | RH | Clonidine 125 micrograms/day | None | 130 | 15 | 5 | 18 | 15 | 38 |
| TS229 | 32.97 | M | RH | Paroxetine | None | 114 | 25 | 15 | 15 | 8 | 38 |
| TS230 | 20.16 | F | RH | None | None | 108 | 18 | 40 | 22 | 13 | 75 |
| TS141 | 12.68 | M | RH | None | None | 129 | 51 | 35 | 13 | 14 | 62 |
| TS207 | 15.28 | M | RH | None | None | 97 | 12 | 0 | 13 | 10 | 23 |
| TS232 | 13.73 | M | RH | Risperidone 3mg/day,<br>Clonidine 35micrograms/day | ADHD | 88 | 13 | 2 | 20 | 19 | 41 |
| TS224 | 19.54 | F | LH | Aripiprazole (25mg), 2mg circadin | ASD | 103 | 14 | 35 | 21 | 21 | 77 |
| TS237 | 14.35 | M | RH | None | ASD, ADHD | 93 | 6 | 38 | 16 | 0 | 54 |
| TS238 | 26.11 | F | RH | Methylphenidate 36mg/day | ADHD | 105 | 16 | 40 | 23 | 21 | 84 |
| TS240 | 35.96 | F | RH | Thyroxine 150mg, Citalopram 20mg,<br>Microgynon 30mg, kalms tablets | None | N/A | 19 | 15 | 22 | 0 | 37 |
| **Average** | 20.36 | - | - | - | - | 104.83 | 21.17 | 26.08 | 18.73 | 12.53 | 58.33 |

WASI: Wechsler's Abbreviated Scale of Intelligence. PUTS: Premonitory Urge for Tics Scale (max score is 32). ADHD: Attention deficit hyperactivity disorder. \*Score taken from YGTSS: YGTSS Global Tic Severity Scale.

### Devices

The visual stimuli were presented on a computer screen (Samsung S23A700D, screen resolution: 1920 × 1080). Participants made responses on an Apple Pro Keyboard M7803 connected to a Mac Pro computer running High Sierra (v 10.13.6) via USB cable. The task was controlled by PsychoPy2 (v1.90.3) script coded using python2 on the Mac Pro.

### Experimental procedure

Participants performed a manual choice Go/NoGo task using the index finger of their dominant hand. Each trial consisted of a fixation cross, presented in the centre of the computer monitor, that disappeared after 1000ms, which was followed by the appearance of the cue stimulus (either a square or a circle) that appeared in the centre of the computer monitor. The task consisted of pressing the response key (space bar) as quickly as possible whenever either a yellow circle or a blue square appeared in the centre of the screen (i.e. the Go condition), or withholding a response whenever a blue circle or a yellow square appeared (i.e. NoGo condition). 70% of the trials were Go trials and 30% of the trials were NoGo trials. The diameter of both stimuli was 5cm. When no response was made, the next trial appeared after 1 second. When a response was made, the next trial began.

Participants were seated approximately 70cm away from the screen. There were 400 trials and a short break was offered every 100 trials. After every break, a reminder of the instructions of the task appeared on the screen. The task did not take longer than 15 minutes. TS participants were told not to suppress their tics.

Before the study commenced, participants completed a set of 20 practice trials. If at least 90% of those trials were performed correctly, the subject proceeded to start the actual study; if less than 90% of those trials were performed correctly, the subject continued to perform another practice of 20 trials until at least 90% of those trials were performed correctly.

### Electroencephalography (EEG) recording

EEG data were recorded from 64 electrodes using a BioSemi Active Two System. Data were recorded with a sampling rate of 1024Hz which was subsequently down-sampled to 128Hz. The impedance of the electrodes was kept under 30μV for all participants. Reference electrodes were placed on the left and right mastoids. Bipolar vertical and horizontal EOG was also recorded.

Data were low-pass filtered at 45Hz and high-pass filtered at 1Hz. Channels showing aberrant behaviour were deleted and noisy channels were interpolated. Automatic Artifact Removal (AAR) was used to remove EOG artifacts using a recursive least squares regression.

Data were standardised (z-scores) to produce data with a mean of 0 and standard deviation to 1. As a result, no baselining of the data was required. Epochs showing abnormal trends or excessive noise were rejected. Independent Component Analysis (ICA) were conducted and artifacts were identified by visual inspection and through the use of the Multiple Artifact Rejection Algorithm (MARA).

### EEG data analyses

Analyses of the EEG data recorded from scalp sensors located over the motor cortex contralateral to the responding hand (i.e., C3 for right-handed participants and C4 for left-handed participants) yielded mean event-related potentials (ERPs) and mean event-related spectral perturbation (ERSP) for correct Go trials. All NoGo trials, and Go trials in which no response was recorded, were discarded from the analyses. In addition, and importantly, all epochs in which the participant was recorded (observed) to be ticcing were discarded from the analyses.

For each correct Go trial, a time window of −1 to 1 seconds was constructed and time-locked to each button press. In each case zero seconds indicated the time of each button press. Statistical analyses were performed separately for mu (8-13Hz) and beta (13-30Hz) frequency responses to test the null hypothesis that the sample data are from a population with mean equal to zero at the 5% significance level. One-sample t-tests, with a test variable of 0, were conducted at each time point correcting for the multiple comparisons using False Discovery Rate (FDR; Benjamini & Hochberg, 1995).

## Results

### Event-related potentials (ERP)

ERP for GO trials for the TS group and healthy controls are presented in Figure 1. Inspection of this figure indicates that for both groups, the ERP increases shortly before movement onset, and there is a sustained increase in ERP during the period of the movement.

**Figure 1.**
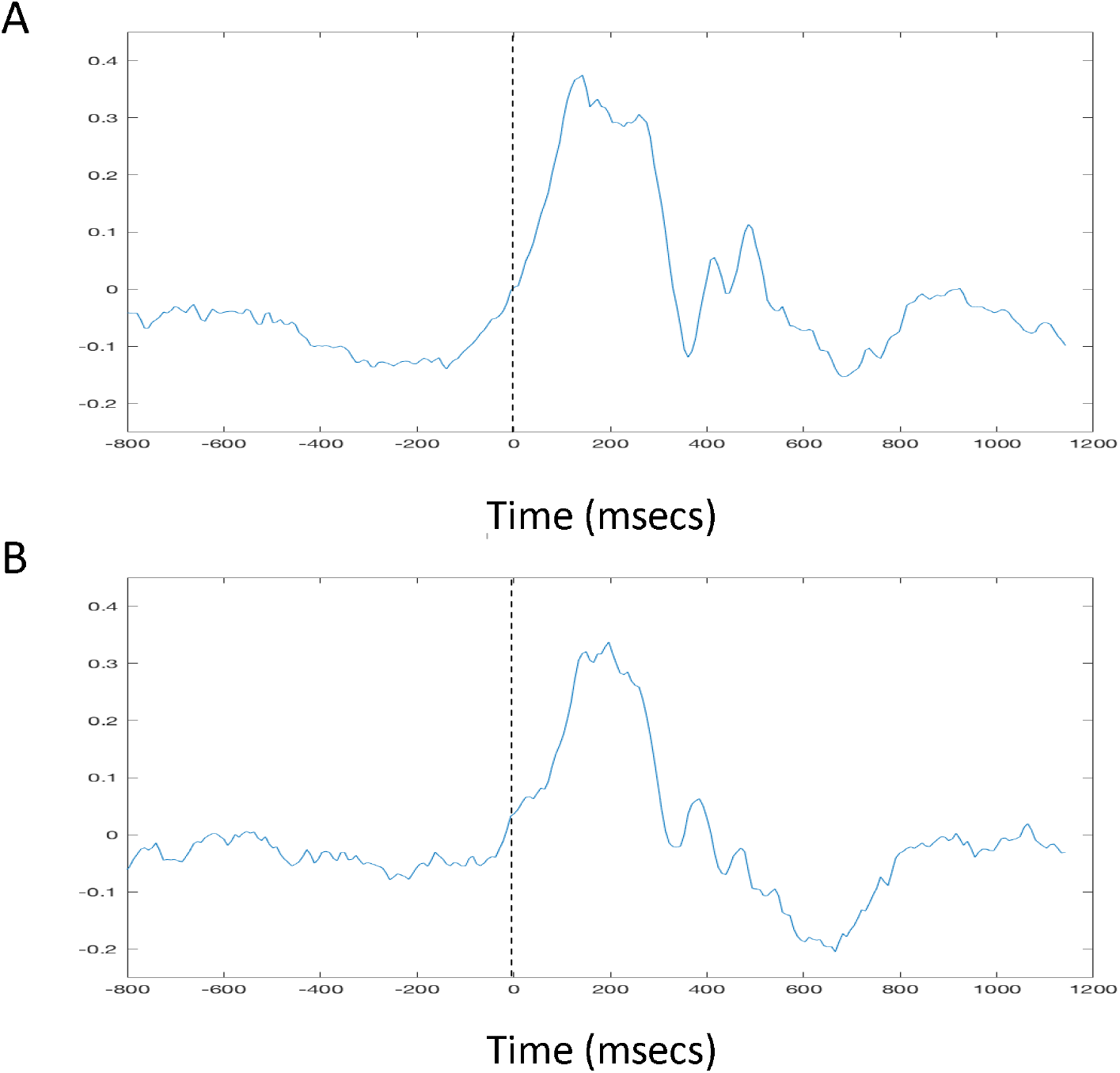
Event-related potentials (ERP) recorded from the scalp over the sensorimotor cortex contralateral to the responding hand. **A**. Illustrates mean ERP data recorded for the healthy control group. B. Illustrates mean ERP data recorded for the TS group. The *vertical* broken black line in each image at time zero on the x-axis indicates the time at which the button press occurred.

### Mu-band ERSP analyses

Time-frequency analyses were conducted for both mu-band and beta-band frequencies. Mu-band time-frequency ERSP data for both the TS group and healthy controls are presented in Figure 2. Inspection of this figure indicates that, for the healthy control group (Figure 2A), there is a strong reduction in mu-band power (desynchronization) that coincides with the movement onset, and statistical analysis using one-sample t-tests (Figure 2C) confirm that the standardised (Z) ERSP values differ from zero (p < .05^FDR-corrected^). By contrast, inspection of Figure 2B suggests that mu-band desynchronization is much weaker in the TS group and occurs some time after movement onset. This is confirmed by the statistical analyses (Figure 2D) which demonstrated that at no time did standardised mu-band power (ERSP) differ significantly from zero.

**Figure 2:**
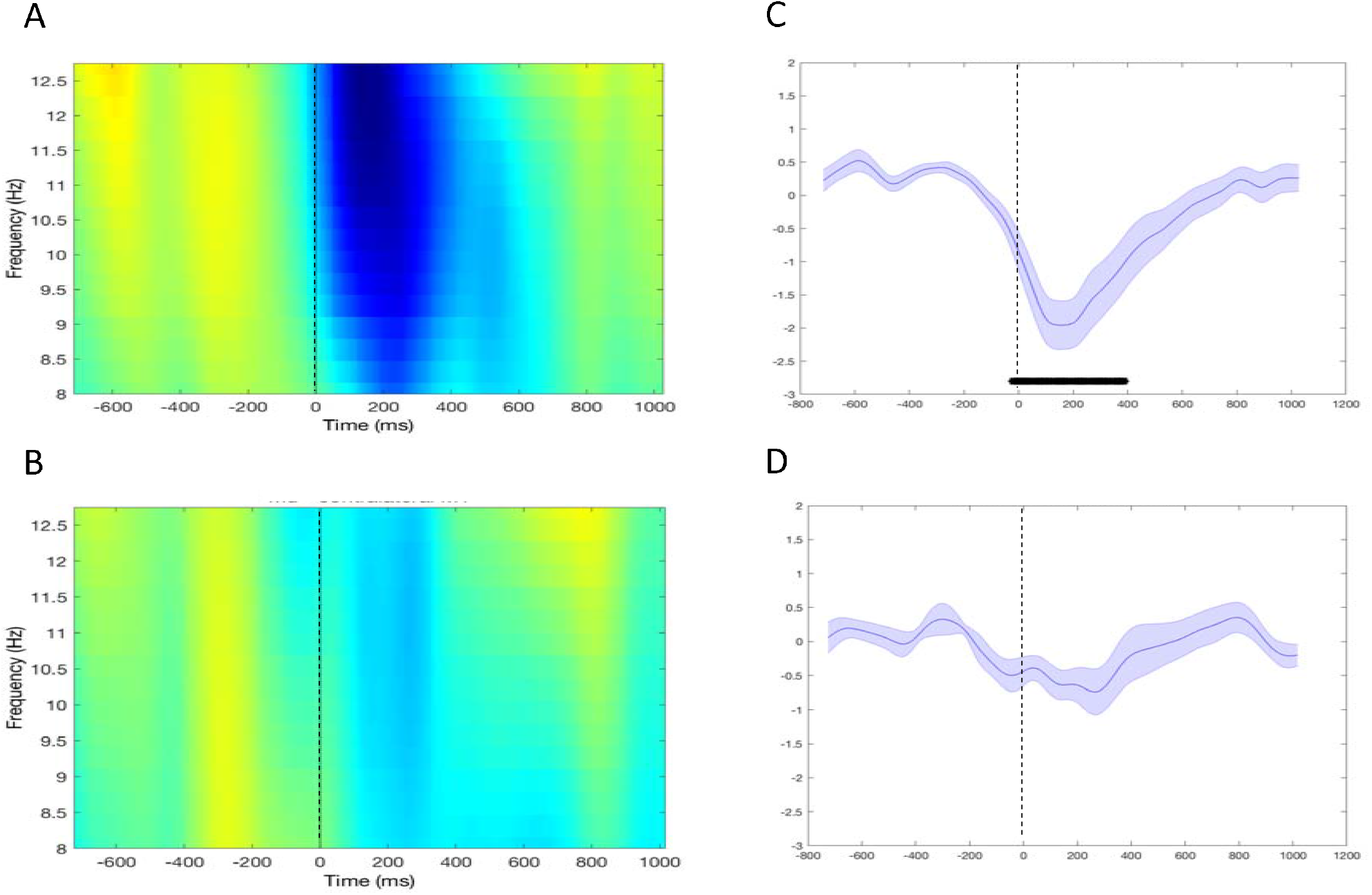
Time-frequency plots showing mu-band frequency power recorded from the scalp over the contralateral sensorimotor area. **A.** Data from healthy controls. **B.** Data from individuals with TS. 12Hz ERSP values recorded from the scalp over the contralateral sensorimotor area. **C.** Data from healthy controls. **D.** Data from individuals with TS. The *vertical* broken black line in each image at time zero on the x-axis indicates the time at which the button press occurred. The black *horizontal* line indicates individual time points at which the ERSP values significantly differed from zero (p < 0.05^FDR-corrected^). The shading in **C** and **D** represents the standard error.

To investigate this further we examined individual event-related desyncronisation (ERD) responses as follows. For each individual we calculated the time of the maximum ERD for that individual. The resultant rank-ordered data are displayed in Figure 3. Further analyses revealed that the mean time of maximum ERD was 242ms (± 129ms) after movement onset for the control group and 227ms (± 215ms) for the TS group. An independent groups t-test revealed that these means did not differ statistically (p > .1). However, it should be noted that the timing of the ERD was considerably more variable in the TS group (Figure 3: coefficient of variation; HC group = 0.53; TS group = 0.95).

**Figure 3.**
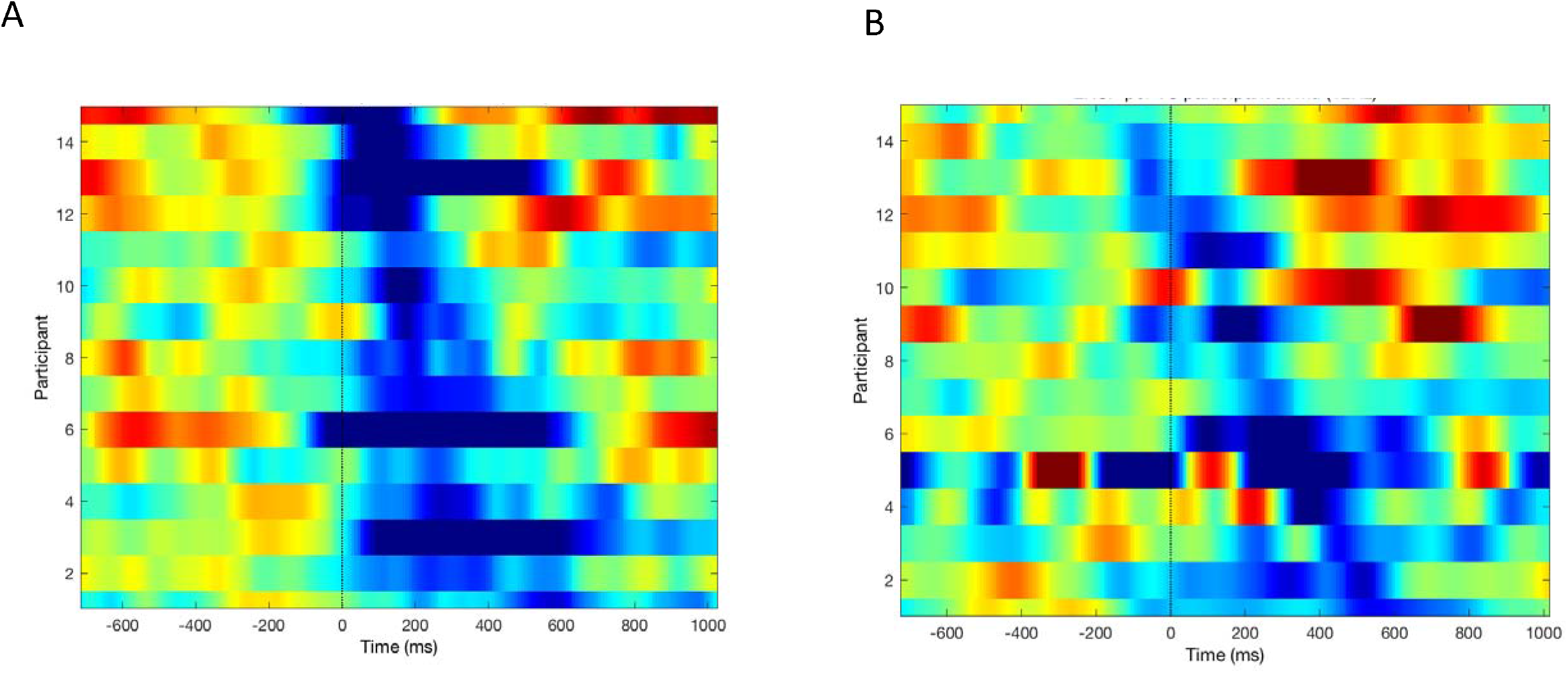
Individual mu-band peak event-related desynchronization (ERD) data was rank-ordered from the earliest (top) occurrence to the latest (bottom) occurrence. **A**. Data from each of the healthy control group is located close to the time of movement onset. **B**. Data from the TS group is much more variable. While the mean time of maximum ERD does not differ across groups the variability is much greater in the TS group (coefficient of variation: HC group = 0.53; TS group = 0.95).

### Beta-band ERSP analyses

Beta-band time-frequency ERSP data for the TS and healthy control groups are presented in Figure 4. Inspection of this figure indicates that, for the healthy control group (Figure 4A), there is a strong reduction in beta-band power (ERD) that begins in advance of movement onset and is sustained for several hundred milliseconds after movement onset. Statistical analysis, using one-sample t-tests (Figure 4C), confirms that the standardised (Z) ERSP values do differ significantly (p < .05^FDR-corrected^) from zero throughout this period. Interestingly, and in contrast to the results for mu-band power, inspection of Figure 4B indicates that beta-band ERD also occurs prior to movement onset TS group, but is sustained for a shorter period of time. This is confirmed by the statistical analyses (Figure 4D).

**Figure 4:**
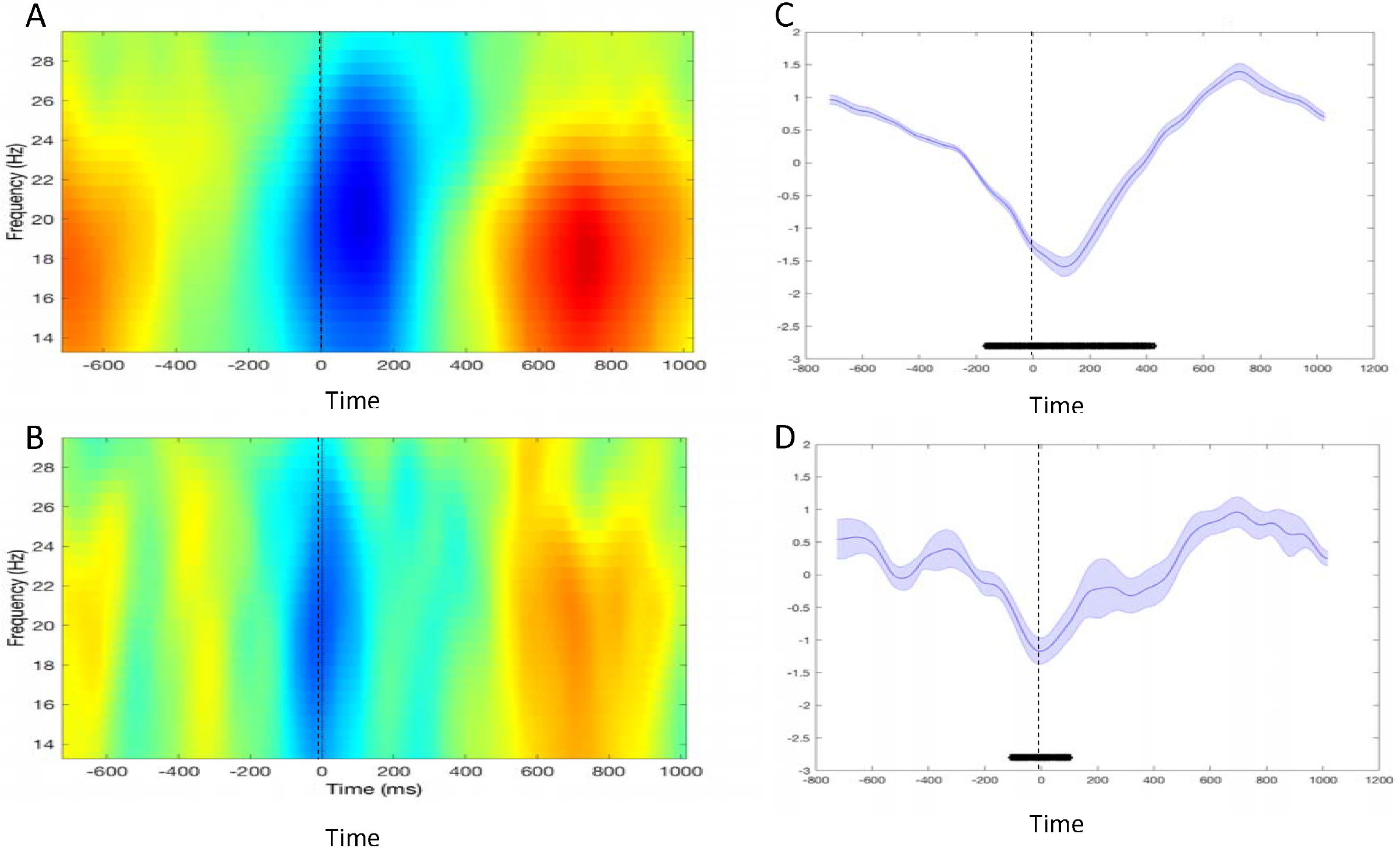
Time-frequency plots showing beta-band frequency power recorded from the scalp over the contralateral sensorimotor area. **A**. Data from healthy controls. **B**. Data from individuals with TS. 19Hz ERSP values recorded from the scalp over the contralateral sensorimotor area. **C**. Data from healthy controls. **D**. Data from individuals with TS. The *vertical* broken black line in each image at time zero on the x-axis indicates the time at which the button press occurred. The black *horizontal* line indicates individual time points at which the ERSP values significantly differed from zero (p < 0.05^FDR-corrected^). The shading in **C** and **D** represents the standard error.

Again, to investigate this further, we examined individual event-related desyncronisation (ERD) responses. For each individual we calculated the time of the maximum ERD for that individual. The resultant rank-ordered data are displayed in Figure 5. Further analyses revealed that the mean time of maximum ERD was 111ms (± 86ms) after movement onset for the control group and 110ms (± 128ms) for the TS group. An independent groups t-test revealed that these means did not differ statistically (p > .1). However, it should be noted that the timing of the ERD was considerably more variable in the TS group (Figure 5: coefficient of variation; HC group = 0.78; TS group = 1.15).

**Figure 5.**
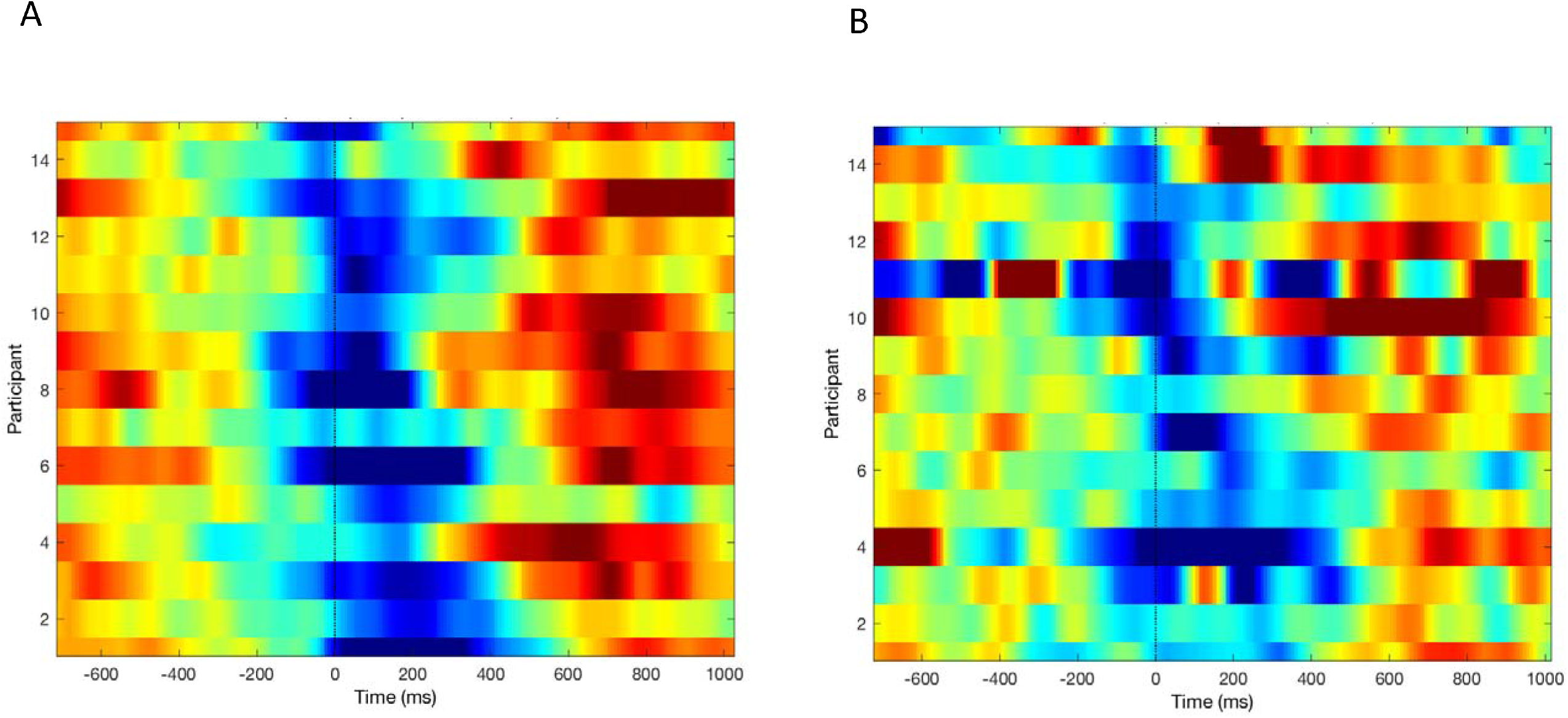
Individual beta-band peak event-related desynchronization (ERD) data was rank-ordered from the earliest (top) occurrence to the latest (bottom) occurrence. **A**. Data from each of the healthy control group is located close to the time of movement onset. **B**. Data from the TS group is much more variable. While the mean time of maximum ERD does not differ across groups the variability is much greater in the TS group (coefficient of variation: HC group = 0.78; TS group = 1.15).

### Between-group analyses of mu and beta power

To test directly whether mu-band and beta-band ERD were significantly different between groups we carried out the following analyses. For each individual, and for each group, we calculated the mean ERSP value for the time period −500ms to +500ms (i..e., 500ms before movement onset to 500ms after movement onset). We then tested this difference in between groups using an independent groups t-test. Separate analyses were conducted for mu and beta power. The analysis revealed that there was no significant between group difference in mean beta band power (control group = −0.31 (± 0.43), TS group = −0.16 (± 0.51); t(28) = −0.88, p = 0.19) but there was a significant difference in mu-band power (control group = −0.72 (± 0.77) decibels, TS group = −0.25 (± 0.55) decibels; t(28) = −1.95, p = 0.03).

## Methods: Study 2. Neural antecedents of motor tics

### Participants

Seventeen individuals (5 female) aged between 12 and 26 years (mean age of 18 years) with a confirmed diagnosis of Tourette Syndrome took part in the study. Participants were recruited from Tourette clinic at the Queen’s Medical Centre, Nottingham and through the UK charity *Tourette’s Action*. All participants (and their parent/guardian where appropriate) provided informed consent to participate in the study which was approved by a University of Nottingham ethical review committee. Participants with co-occurring conditions were not excluded. One participant’s data had to be rejected due to there being no intervals between their tics. The 16 participants that remained included 4 females aged between 12 - 23 years (mean age of 17.4 years). Of the sixteen subjects, 5 were currently taking medication. Details from participants can be found in Table 2.

**Table 2.** Characteristics of TS participants in Study

| TS ID | Age | Gender | Medication | Comorbidity | WASI | PUTS | Impairment* | Motor* | Vocal* | Motor+Vocal* | Global* |
| --- | --- | --- | --- | --- | --- | --- | --- | --- | --- | --- | --- |
| TS030 | 21.1 | M | None | None | 103 | 5 | 10 | 13 | 11 | 24 | 34 |
| TS061 | 14.8 | M | Clonidine (75mg/day) | None | 99 | 14 | 2 | 19 | 16 | 35 | 37 |
| TS062 | 22.7 | M | None | None | 131 | 22 | 35 | 22 | 22 | 44 | 79 |
| TS075 | 20.2 | M | None | None | - | 17 | 15 | 12 | 10 | 22 | 37 |
| TS077 | 23.1 | M | None | ADHD | 91 | 8 | 10 | 17 | 18 | 35 | 45 |
| TS139 | 14.4 | M | None | None | 100 | 5 | 25 | 12 | 0 | 12 | 37 |
| TS141 | 12.7 | M | None | None | 129 | 10 | 35 | 13 | 14 | 27 | 62 |
| TS171 | 18 | F | None | None | 101 | 9 | 8 | 9 | 6 | 15 | 23 |
| TS176 | 12.5 | M | None | None | 107 | 4 | 17 | 12 | 13 | 25 | 42 |
| TS178 | 18.2 | F | Aripiprazole 10mg/day | None | 110 | 25 | 30 | 17 | 20 | 37 | 67 |
| TS182 | 17.4 | F | None | None | 85 | 24 | 15 | 17 | 9 | 26 | 41 |
| TS183 | 12.5 | M | None | None | 139 | 11 | 35 | 18 | 18 | 36 | 71 |
| TS184 | 19.4 | M | Sulpiride 400mg | None | 82 | 30 | 50 | 21 | 21 | 42 | 92 |
| TS189 | 12.4 | M | Depakote 750mg, Clonidine 0.15mg, Quetiapine 200mg | None | 99 | 21 | 30 | 18 | 18 | 36 | 66 |
| TS190 | 23.5 | M | None | None | 114 | 30 | 40 | 18 | 10 | 28 | 68 |
| TS191 | 15.9 | F | None | ADHD, OCD | 113 | 23 | 30 | 19 | 2 | 21 | 51 |
| **Average** | **17.4** | **-** | **-** | **-** | **106.86** | **16.12** | **24.18** | **16.06** | **13** | **29.06** | **53.25** |

WASI: Wechsler's Abbreviated Scale of Intelligence. PUTS: Premonitory Urge for Tics Scale (max score is 32). ADHD: Attention deficit hyperactivity disorder. OCD: Obsessive compulsive disorder. \*Score taken from YGTSS: YGTSS Global Tic Severity Scale.

### EEG data collection

EEG data collection was identical to that described for Study 1. Two EMG electrodes were placed over the *orbicularis oris* in order to record the onset of tics involving mouth and nose movements. Time-windows of −1.2 to 0.5 seconds, time-locked to the tic onset were extracted. The average number of epochs across participants was 42.

### Experimental procedure

During EEG recording all participants eye and facial movements were video recorded for off-line analysis of eye and facial tics. At the start of the video recording participants were asked to sit in a relaxed posture and were instructed not to suppress their tics. Testing took ~28 minutes with three small breaks every 7 minutes.

### Selection and marking of tic events

EEG data was synchronised with the video recorded during the testing. Off-line video analysis was conducted using QuickTime 7 Pro player allowing frame-by-frame analysis (maximum error range 20ms). Once the EEG recording was synchronised with the video, all tics that occurred in a period at least 1 second from the last recorded tic, or from a voluntary movement, were marked in the EEG data. This process was performed twice for each participant to ensure reliability of identifying tic onsets. As a result, only tics that were preceded by at least one second of resting state (i.e., no movement observed), and could be clearly identified in the EEG data were marked. Because of these conservative requirements, some tics, particularly those that occurred in a sequence separated by less than a second, may not have been identified as individual events.

### EEG data Analysis

As outlined above, analyses of the EEG data were conducted to obtain ERP and ERSP for the electrodes located on the scalp over the sensorimotor cortex (i.e., average ERP or ERSP from electrodes C3, C1, C2, C4).

## Results

The average event-related potential (ERP), before and after tic onset, is presented in Figure 6A. Inspection of this figure clearly indicates that the ERP at tic onset does not exhibit the voltage increase observed for voluntary movements executed by healthy individuals (Figure 1A) and individuals with TS (Figure 1B).

**Figure 6.**
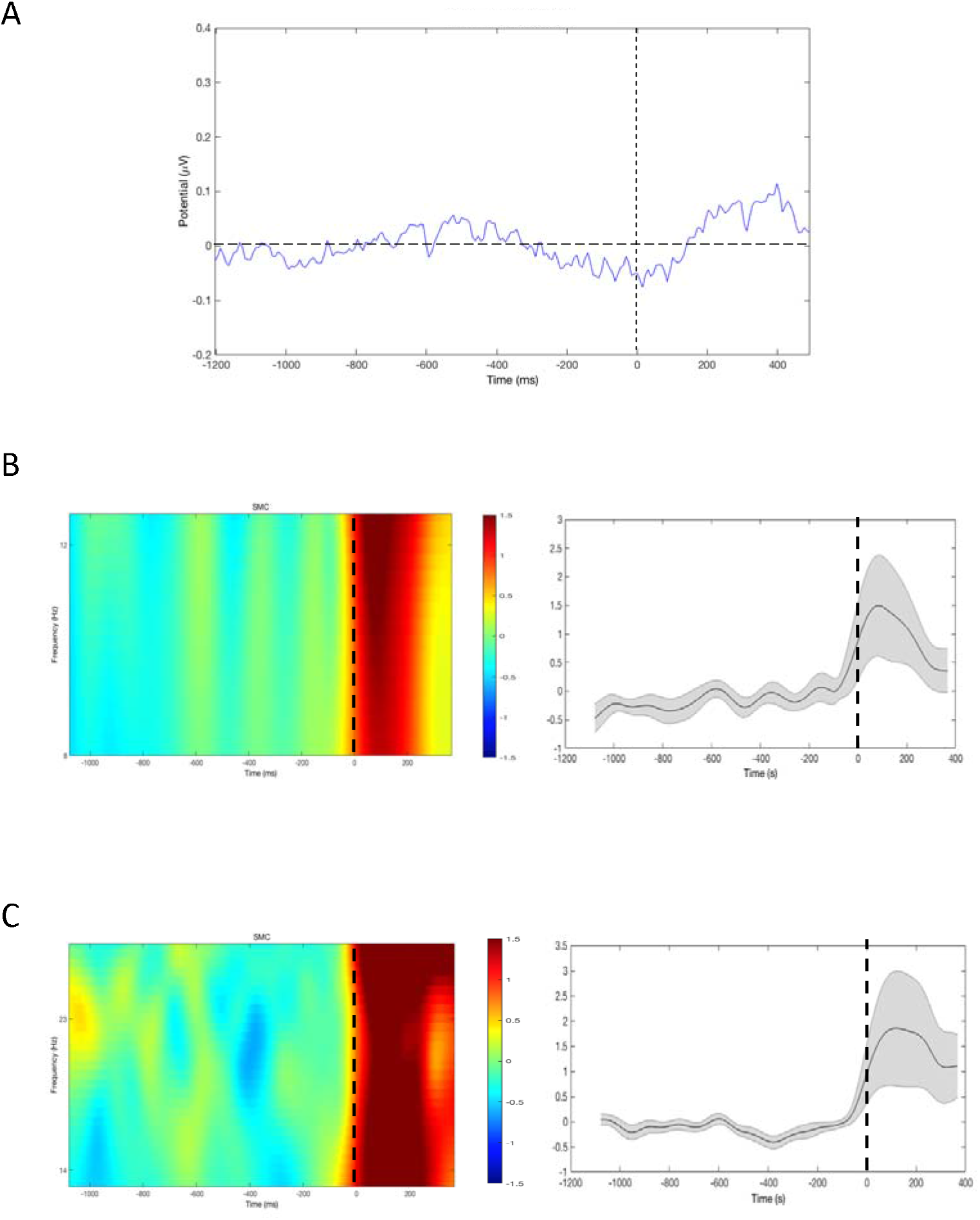
A. Average ERP from the average electrodes on the SMC. The broken vertical black line indicates the start of the tic. B. Left panel: Time-frequency plots in mu-band (8-13Hz) for the average electrodes on the SMC. Right panel: Values of power for the average electrodes on the SMC. C. Left panel: Time-frequency plots in beta-band (13-30Hz) for the average electrodes on the SMC. Values of power for the average electrodes on the SMC in mu-band (8-13Hz). Right panel: Values of power for the average electrodes on the SMC in beta-band (13-30Hz). 0 indicates the onset of the tic.

To investigate this further, we conducted further time-frequency analyses for mu-band (Figure 6B) and beta-band (Figure 6C) power (ERSP), time-locked to the onset of tics (0ms). These time-frequency analyses demonstrate that, in contrast to voluntary movements, there was no substantive desynchronization of power immediate prior to, or at the time of tic onset, for either mu or beta rhythm (see Figure 4). Figure 6 does appear to show an apparent increase in ERSP across multiple band widths, however it is important to recognise these this is unlikely to reflect an event-related increase in power, but instead almost certainly reflects muscle activity (artefact) generated as a consequence of tic-related facial/head movement, and should therefore be discounted and will not be discussed.

For consistency with the analyses reported for Study 1, we examined individual event-related desyncronisation (ERD) responses prior to the onset of each tic. For each individual, and for each trial, we calculated the time of the maximum ERD. The resultant data were rank-ordered and are displayed in Figure 7. Inspection of this figure confirms that there is no evidence for either a mu-band or beta-band desynchronization immediately preceding, or coinciding with, tic onset.

**Figure 7.**
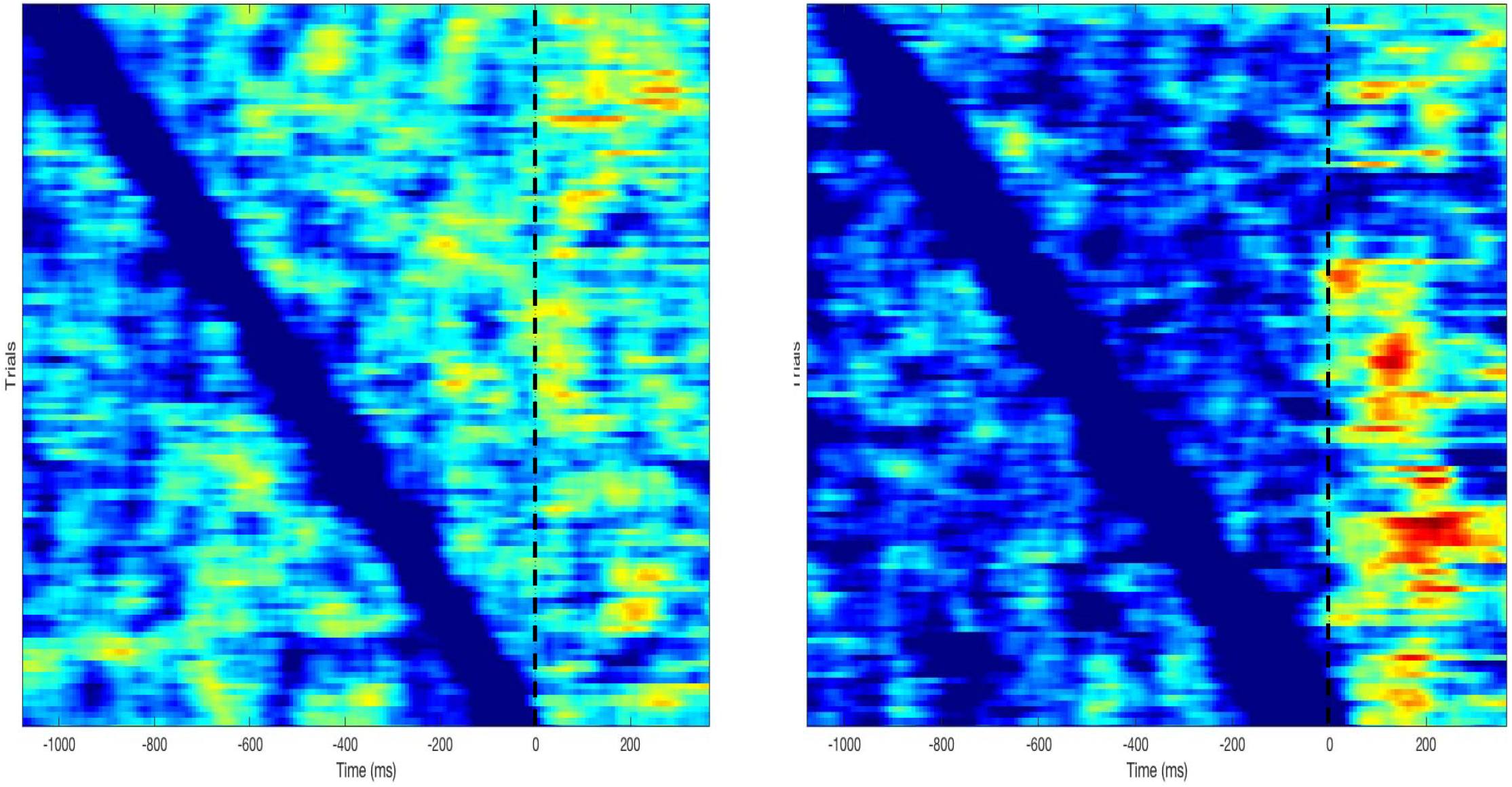
Averaged event-related desynchronization (ERD) peaks, recorded for the electrodes located over the sensorimotor cortex, for individual trials rank-ordered according to the time of occurrence of the earliest (top) to latest (bottom) peak ERD prior to tic onset. Left panel: data for mu-band ERD. Right panel data for beta-band ERD. The broken black vertical line centred at 0ms indicated the time of tic onset in each case.

## Discussion

We examined the neural antecedents of voluntary movements and of motor tics using EEG recording techniques. Our results can be summarised as follows. First, we observed that, for voluntary movements, event-related electrical potentials (ERP) recorded from the scalp over the sensorimotor cortex contralateral to the responding hand were highly similar for individuals with TS compared to typically developing individuals. Second, time-frequency analysis of movement-related beta-band oscillation power also revealed that both the TS group and controls exhibited an event-related desynchronization (ERD) that commenced prior to movement onset. However, a similar time-frequency analysis of mu-band oscillation power revealed that, whereas the healthy control group exhibited a clear mu-band ERD that commenced at movement onset, this was not observed for the TS group. Third, and in our view importantly, investigation of the temporal variability of the onset of peak ERD for both mu and beta oscillations demonstrated that while the average time of peak ERD did not differ between groups, the timing of peak ERDs was considerably more variable for the TS group. This will be discussed below. Finally, analysis of ERP prior to, and at the time of, tic movements did not reveal the positive-going increase in ERP amplitude observed for voluntary movements. Furthermore, time-frequency analyses of both mu-band and beta-band power confirmed that there was no evidence of any ERD immediately prior to the onset of tics.

### Absence of beta and mu band oscillations prior to tics in TS

Neural oscillations of the brain’s electromagnetic activity reflect the synchronised firing of populations of neurons, and it is known that GABA-mediated interneurons play a critical role in co-ordinating the synchronised activity of populations of pyramidal neurons that give rise to brain oscillations (Schnitzler & Gross, 2005). Two frequency bands are particularly relevant to the occurrence of tics in TS: Mu (8-12Hz) and Beta (13-30Hz); which have long been associated with sensorimotor function (Armstrong, Sale, Cunnington, 2018); become de-synchronised when a movement is initiated (Schnitzler & Gross, 2005); and, have been linked to maintaining the current motor set (Engel & Fries, 2010). However, a more recent proposal is that beta-band oscillation power reflects an internal estimate of the likelihood of the need for a novel voluntary action, and effectively signals the currently level of motor readiness (Jenkinson and Brown, 2011). Furthermore, it is argued that beta-band oscillations are linked to net dopamine levels within key sites in cortical-striatal-thalamic-cortical motor networks (Jenkinson and Brown, 2011). Within this context it is of interest to note that Parkinson’s disease - a disorder that is associated with a poverty of spontaneous movements (Akinesia), slowed movements (Bradykinesia), and is caused by a loss of dopamine cells within the substantia nigra – is associated with abnormally increased power in beta-band oscillations, that is reduced by dopamine medication and deep-brain stimulation. Moreover, the extent to which bradykinesia improves following drug treatment is related to the extent to which beta-band oscillation power is reduced within the cortical-striatal-thalamic-cortical motor circuit (Jenkinson and Brown, 2011). By contrast, Tourette syndrome – a disorder that is associated with the occurrence of involuntary movements and vocalisations (tics), and is associated with increased dopamine activity within the cortical-striatal-thalamic-cortical motor circuit (Buse, Schoenefeld, Münchau, Roessner, 2012) – is shown in the current study to be associated with an absence of beta and mu-band desynchronization ahead of the occurrence of motor tics. While this finding is consistent with the results of previous EEG studies that demonstrate an absence of movement-related ERP prior to tics in TS, it is nonetheless rather puzzling as to why no movement-related EEG signals (i.e., ERD) are observed immediately prior to, or during, tic onset, as functional magnetic resonance imaging (fMRI) studies of this issue have clearly demonstrated increased brain activity in sensorimotor cortex both immediately before tic onset and at tic onset. Specifically, an early study examined brain activity occurring 2 seconds before tic onset and also at tic onset, and found widespread increases in activation in paralimbic areas such as anterior cingulate cortex and insular, the supplementary motor area and parietal operculum 2 seconds prior to tic, whereas during tic execution activity was observed in primarily sensorimotor cortex and the cerebellum (Bohlhalter, Goldfine, Matteson, Garraux, Hanakawa, Kansaku, Wurzman, Hallett, 2006). Similarly, a more recent study with a far larger sample size, largely confirmed this finding and reported increased activity within the sensorimotor cortex both 1 second before, and at, tic onset (Neuner, Werner, Arrubla, Stöcker, Ehlen, Wegener, Schneider, Shah, 2014). These studies confirm that tic generation is associated with a set of brain areas that include, in particular, the insula and cingulate cortices, the cerebellum, and basal ganglia nuclei, which may indicate that the generation of tics in TS may involve brain activity that is not typically observed for volitional movements. This is supported by the striatal disinhibition animal model of TS. In this model, tic-like movements are produced following localised injections of a GABA-antagonist into the striatum of the animal (Worbe et al., 2009; Worbe et al., 2013; Bronfeld et al., 2013; McCairn et al., 2013). McCairn and colleagues conducted multisite, multielectrode electrophysiological recordings of single-unit activity and local field potentials (LFPs) from a number of brain areas in non-human primates, including: the cerebellum; basal ganglia; and primary motor cortex, while the animal exhibited tic-like movements. A key finding from their study was that, while changes in LFPs preceded the occurrence of tic-like movements in many of the brain areas recorded from with different latencies, the occurrence of tic-like movements was most closely associated with LFPs in the primary motor cortex and cerebellum (McCairn et al., 2013). The authors propose that while striatal disinhibition might be a trigger for tic-like movements, the primary motor cortex, and in particular the cerebellum, may act as a gate to release overt tic-like movements (McCairn et al., 2013).

One difference between volitional movements and tics is that tics are most often preceded by a premonitory sensory/urge phenomena (PU) which are consciously experienced as a strong urge for motor discharge and have been thought of as a driver for tics. Importantly, these PU have been particularly linked to the insular cortex, with the mid-cingulate cortical region associated with the triggering of actions in response to PU (Jackson et al., 2011). In this context, it is important to note that brain activity within the mid cingulate cortex increases immediately prior to the execution of tics (Bohlhalter et al., 2006; Neuner et al., 2014) and that there are alterations in grey matter (GM) volume throughout mid and anterior cingulate cortex that are correlated with clinical measures of tic severity (O’Neill, Piacentini & Peterson, 2019). Furthermore, direct electrical stimulation of the mid-cingulate cortex is sufficient to induce movements, including goal-directed actions, with no evidence that electrical stimulation of this region induces a phenomenological experience of an ‘urge-to-move’ (Caruana, Gerbella, Avanzini, Gozzo, Pelliccia, Mai, Abdollahi, Cardinale, Sartori, Lo Russo, Rizzolatti, 2018; Trevisi, Eickhoff, Chowdhury, Jha, Rodionov, Nowell, Miserocchi, McEvoy, Nachev, Diehl, 2018). Finally, recent brain imaging studies investigating differences in seed-based structural covariance networks (SCNs) in individuals with TS compared to matched controls have demonstrated that the SCNs linked to the insula (Jackson, Loayza, Crighton, Sigurdsson, Dyke, Jackson, 2020) and mid cingulate cortex (Jackson, Sigurdsson, Dyke, Condon, Jackson, 2020), and the motor cerebellum (Sigurdsson, Jackson, Jolley, Mitchell, Jackson, 2020) all have substantially different patterns of structural covariance compared to typically developing individuals. Importantly, each study indicates that structural covariance is substantially increased between the insula, mid cingulate in individuals with TS relative to controls. Taken together, these studies lend support to the proposal that the contribution of paralimbic, sub-cortical, and cerebellar regions may play a far greater role in the generation of tics in TS, than they necessarily do in the generation of volitional movements.

### Difference between mu and beta band oscillations during voluntary movements in TS

As noted above, in the current study we observed that the ERP recorded from the scalp over the sensorimotor cortex prior to volitional movements was highly similar for both individuals with TS and typically developing individuals, and the magnitude of the beta-band ERD observed for the TS group did not differ from that of the matched control group. By contrast, analysis of mu-band power revealed that while the healthy control group exhibited a clear mu-band ERD that commenced at movement onset, this was not observed for the TS group, and mu power for the per-movement period was significantly different between the groups. Below we discuss two potential explanations for this observation, both of which can be linked to altered physiological inhibition in TS.

First, it has been suggested that TS is associated with increased levels of sensorimotor noise that accompany tics and result in it being more difficult for individuals with TS to distinguish tics from volitional movements (Ganos et al., 2015). Evidence in support of this proposal comes from a recent study that examined forward model updating in young adults with TS (Kim, Jackson, Dyke, Jackson, 2019). Thus, it may be that sensorimotor processes, including the timing of movement-related brain oscillations, are less precise in individuals with TS; perhaps as a consequence of alterations in GABA-mediated inhibitory processes which are known to play a key role co-ordinating the synchronised activity of populations of pyramidal neurons that give rise to brain oscillations (Schnitzler & Gross, 2005; Prokic et al., 2019), and are reported to be abnormal in individuals with TS (Gilbert et al., 2004; Kalanithi et al., 2005; Orth & Rothwell, 2009; Lerner et al., 2012; Puts et al., 2015). Specifically, there are now multiple lines of evidence to indicate that TS is associated with altered GABA function within cortical-striatal-thalamo-cortical motor networks. Post-mortem investigations of TS have demonstrated substantial decreases (~50%) in the number of GABAergic interneurons found within the striatum (Kalanithi et al., 2005), and Positron Emission Tomography (PET) imaging has revealed widespread alterations in GABA_A_ receptor binding in TS, in particular within the striatum, thalamus, insula, and primary somatosensory cortex (Lerner et al., 2012). In addition, Transcranial Magnetic Stimulation (TMS) studies of physiological inhibition within the primary motor cortex have demonstrated reduced GABA_A_ receptor dependent short intra-cortical inhibition (Gilbert et al., 2004; Orth & Rothwell, 2009) and reduced interhemispheric inhibition (IHI) in TS (Bäumer et al., 2010), and a Magnetic Resonance Spectroscopy (MRS) study has reported decreased concentrations of GABA within the primary sensorimotor cortex in children with TS (Puts et al., 2015). In the current study, consistent with the proposal for increased sensorimotor noise in TS, we observed in Study 1 that, while the mean timing of movement-related desynchronization (ERD) did not differ from controls in the TS group, the *variability* in the timing of the peak ERD relative to movement onset was substantially larger in the TS group for both mu and beta band oscillations.

While both mu and beta band oscillations are associated with the initiation of voluntary movements, it is important to recognise that they may nonetheless each index different sensorimotor functions. As noted above, beta-band oscillations are thought to play a role in movement preparation by reflecting an internal estimate of the likelihood of the need for a novel voluntary action and signalling the current state of motor readiness (Jenkinson and Brown, 2011). As such, it is likely that beta-band desynchronization is associated with the disinhibition (activation) of movement-related neuronal populations. By contrast, mu-band oscillations are thought to play a key role in inhibiting neuronal populations that might otherwise interfere with movement selection (Jensen & Mazaheri, 2010; Brinkman, Stolk, Dijkerman, de Lange, Toni, 2014; Brinkman, Stolk, Marshall, Esterer, Sharp, Dijkerman, de Lange, Toni, 2016).

Previous studies have reported that individuals with TS exhibit reduced cortical excitability in the motor cortex during movement preparation compared to healthy controls (Draper, Jude, Jackson, & Jackson, 2015; Heise et al., 2010; Jackson et al., 2013); and fMRI studies have also shown a decreased BOLD signal in the motor cortex during execution of movement in individuals with TS compared to healthy controls (Jackson et al., 2011; Roessner et al., 2012; Thomalla et al., 2014). In the current study we found that when executing voluntary movements, the mean amplitude of movement-related beta-band ERDs, while somewhat reduced in the TS group, did not differ significantly to that of the matched health control group. By contrast, we observed that while the control group exhibited a significant movement-related mu-band ERD, this was not observed for the TS group, and the magnitude of the ERD was significantly different across the groups. These findings suggest that while processes involves in the disinhibition of movement-related populations within the sensorimotor cortex may be working effectively in individuals with TS, those neural mechanisms involved in suppressing processes that might otherwise compete with movement generation are working less efficiently in TS. This finding is entirely consistent with the pathophysiology of TS which has been linked with reduced physiological inhibition within the cortico-striatal-thalamo-cortical motor circuit (Albin & Mink, 2006). Specifically, it is consistent with the finding that short interval intracortical inhibition (SICI), a phenomenon that is thought to be dependent upon GABA interneurons, and mediated by GABA_A_ receptors, is significantly reduced in individuals with TS (Gilbert et al., 2004; Orth & Rothwell, 2009). It is also consistent with the finding that somatosensory function (i.e., tactile detection and adaptation tasks) was impaired, and GABA concentrations within primary sensorimotor cortex reduced, in children with TS compared to a matched control group (Puts et al., 2015). Furthermore, this study showed that sensorimotor GABA concentrations were associated with both the severity of motor tics and altered somatosensory function in the children with TS. Together these studies indicate that individuals with TS may exhibit an impairment in physiological inhibition mechanisms that would likely impact on their ability to effectively inhibit competing neuronal populations during movement preparation.

In summary, the current study has confirmed that movement-related mu and beta band oscillations, that have been linked to the initiation of volitional movements, are not observed prior to tics in individuals with TS. We interpret this effect as reflecting the greater involvement of a network of brain areas, including the insular and cingulate cortices, basal ganglia nuclei, and the cerebellum, in the generation of tics in TS. By contrast, beta-band desynchronization is demonstrated to occur when individuals with TS initiate voluntary movements, but, in contrast to healthy controls, desynchronization of mu-band oscillations is not observed during the execution of voluntary movements for individuals with TS. We interpret this finding as reflecting a dysfunction of physiological inhibition in TS, thereby contributing to an impaired ability to suppress neuronal populations that may compete with movement preparation processes.

## Acknowledgements

This work was supported by Tourettes Action (UK) and the NIHR Nottingham Biomedical Research Centre. The views expressed are those of the authors and not necessarily those of the NHS, the NIHR or the Department of Health. We thank Tourettes Action (UK) for assisting with participant recruitment.

